# Direction of gait asymmetry following stroke determines acute response to locomotor task

**DOI:** 10.1101/588103

**Authors:** Virginia L Little, Lindsay A Perry, Mae WV Mercado, Steven A Kautz, Carolynn Patten

## Abstract

**Background:** Given the prevalence of gait dysfunction following stroke, walking recovery is a primary goal of rehabilitation. However, current gait rehabilitation approaches fail to demonstrate consistent benefits. Furthermore, asymmetry is a prominent feature of gait dysfunction following stroke. Differential patterns of gait asymmetry may respond differently to gait training parameters.

**Objective:** The purpose of this study was to determine whether differential responses to locomotor task condition occur on the basis of direction of step length asymmetry (Symmetrical, NP_short_, P_short_) observed during overground walking.

**Methods:** Participants first walked overground at their self-selected walking speed. Overground data were compared against three task conditions all tested during treadmill walking: self-selected speed with 0% body weight support (TM); self-selected speed with 30% body weight support (BWS); and fastest comfortable speed with 30% body weight support and nonparetic leg guidance (Guidance_NP_). Our primary outcomes were: step length, single limb support duration, and stride length.

**Results:** We identified differences in the response to locomotor task conditions for each step length asymmetry subgroup. Guidance_NP_ induced an acute spatial symmetry only in the NP_short_ group and temporal symmetry in the Symmetrical group.

**Conclusions:** Task conditions consistent with locomotor training do not produce uniform effects across subpatterns of gait asymmetry. We identified differential responses to locomotor task conditions between groups with distinct asymmetry patterns, suggesting these subgroups may require unique intervention strategies. Despite group differences in asymmetry characteristics, improvements in symmetry noted in the Symmetrical and NP_short_ groups were driven by changes in both the paretic and nonparetic limbs.

## Introduction

Nearly 80% of stroke survivors regain the ability to walk, yet most are left with persistent gait dysfunction.^1^ Indeed, stroke often results in an asymmetric, functionally demanding, and metabolically inefficient walking pattern.^2,3^ As a result, recovery of walking function is one of the most frequently articulated rehabilitation goals among stroke survivors.^4^ Regardless, only half of people undergoing gait rehabilitation following stroke demonstrate improvements in their walking.^5,6^

Task-specific interventions have become favored as a means to facilitate neuromotor recovery following stroke.^7–11^ One such example, locomotor training (LT), is a contemporary task-specific intervention founded in the rationale that applying appropriate sensory inputs^†^ drives motor recovery. These sensory inputs are arguably facilitated by using a permissive environment which typically includes a treadmill, partial body weight support, and manual assistance.^7^ However, study results regarding the benefits of LT have been equivocal, at best.^5,12,13^

Asymmetry is a prominent feature of gait dysfunction following stroke. While gait asymmetry is broadly recognized, the best approach to mediate these effects has yet to be established.^14–19^ Prior work suggests each seemingly subtle decision made in the course of training, including treadmill speed, handrail hold, and harness support, can impact gait symmetry following stroke.^16,20–22^ Indeed, both handrail hold and partial body weight support (BWS) are argued to provide stability through added sensory cues, reducing the task demands of walking and normalizing the gait pattern toward symmetry.^16^ However, one possibility as to why LT has failed to demonstrate consistent effects is that the LT paradigm was not explicitly designed to remediate gait asymmetry.

Compounding this issue, different patterns of asymmetry have been noted among stroke survivors.^23–26^ Yet, much work arguing for interventions to reduce gait asymmetry often ignores the directionality of asymmetry, reporting instead only the magnitude of asymmetry.^13,26,27^ Ignoring the directionality of asymmetry limits the interpretative power of a finding of reduced asymmetry. Further, individuals with different gait asymmetry patterns might respond differently to the treatment, thus, positive, or desired effects can be cancelled out when combined with absent or negative responses. There is, therefore, a need to understand how people with different asymmetry patterns respond to locomotor training parameters.

Additionally, gait asymmetries can be quantified in numerous ways, but generally fall into two broad categories: spatial and temporal.^19,25,26,28–30^ Spatial asymmetries are commonly quantified by relating the step length of the paretic leg to the nonparetic, whereas temporal asymmetries quantify the relationship between swing time or single limb support time.^23,26,28^

Here we studied how the three key components of LT influence the biomechanics of gait post stroke and whether the effects differ by spatial asymmetry pattern. Specifically, we aimed to determine how changes in task condition including: treadmill walking (TM), BWS, and manual guidance of the nonparetic limb (Guidance_NP_) influence the spatiotemporal parameters of walking post stroke. To determine the differential effects of task condition on spatial asymmetry, we investigated these changes relative to three spatial asymmetry patterns: 1) symmetrical step lengths (Symmetrical), 2) shorter paretic step length than nonparetic (P_short_), and 3) shorter nonparetic step length than paretic (NP_short_).

## Methods

### Participants

We studied 39 individuals with chronic, post-stroke hemiparesis, able to walk independently at least 10 meters with an ankle foot orthosis (AFO) or assistive device. Participants were excluded if they demonstrated any of the following: severe perceptual or cognitive deficits, significant lower extremity contractures or joint pain, cardiovascular impairments contraindicative of walking, body weight exceeding 300 pounds, pathological fracture, or profound sensory deficits. The University of Florida Health Science Center Institutional Review Board (#160-2008) approved all procedures described herein and all participants provided written informed consent prior to participation.

### Protocol Overview

Data were collected in two experimental sessions. At the first session participants were tested with: 1) clinical metrics to assess motor impairment, gait and balance function and 2) a GAITRite^®^ Electronic Walkway (CIR systems Inc., Sparta, NJ) to obtain spatiotemporal gait parameters while walking overground. Participants were familiarized with the partial BWS environment that included: walking on an instrumented split belt treadmill (Bertec, Columbus, OH) while wearing a modified mountain climbing harness (Robertson Mountaineering, Henderson, NV) with partial BWS (Therastride, St. Louis MO). The second session involved motion analysis while walking on the treadmill under three locomotor task conditions detailed below.

### Session 1

#### GaitRite^®^

During overground walking all participants wore comfortable clothing and walking shoes and were permitted to use an assistive device if needed (n= 3), but not an AFO. If needed, an aircast (DJO, Vista, CA) was used to provide medial-lateral ankle stability (n=5). Participants walked twice over the GAITRite^®^ mat; the data from these two trials were averaged to obtain self-selected walking speed. The following outcome measures were extracted using GAITRite^®^ Software (Version 3.9, Sparta, NJ): step length, stride length, single-limb-support percent (SLS%), and gait speed.

#### Step Length Asymmetry Categorization

To determine the presence and direction of step length asymmetry during overground walking, we calculated a paretic step ratio (PSR) which quantifies the proportional contribution of the paretic step to the stride length.^24^ We categorized our sample according to PSR values as follows: 1) *symmetrical step lengths* (Symmetrical; 0.475 ≤ PSR ≤ 0.525), 2) *paretic step length shorter than nonparetic* (P_short_; PSR < 0.475), and 3) *nonparetic step length shorter than paretic* (NP_short_; PSR > 0.525).^25^

#### Quantifying Temporal Asymmetry

To provide descriptive statistics for temporal asymmetry, we calculated a temporal symmetry index (TSI) similar to our PSR calculation used for spatial symmetry with the following equation: TSI =SLS%_paretic_ / (SLS%_paretic_ + SLS%_nonparetic_), where SLS%_paretic_ and SLS%_nonparetic_ are the portions of the gait cycle spent in single limb support on the paretic and nonparetic limbs, respectively. A TSI value of 0.5 represents temporal symmetry.

#### Familiarization: Partial Body Weight Support and Limb Guidance

To become familiar with the LT paradigm, each participant walked on the treadmill for short bouts in combinations of their preferred self-selected and fastest comfortable speeds and 0% and 30% BWS. All participants were then provided manual assistance while walking at their fastest comfortable speed with 30% BWS. Assistance was provided at the nonparetic foot to promote increased step length and normalization of step timing. Each participant performed up to three 5-minute bouts with standing or seated rest breaks provided between bouts. Vital signs, including blood pressure and heart rate, were monitored at baseline and during rest periods.

### Session 2

Instrumented gait data including kinematics and kinetics were acquired using 12 infrared cameras (Vicon MX, Vicon Motion Systems Ltd., Oxford, UK; sampling frequency: 200Hz) and a modified Helen Hayes marker set (41 single reflective markers and 11 rigid clusters) as participants walked on an instrumented split-belt treadmill (Bertec, Columbus, OH).

#### Locomotor Task Conditions

Three treadmill walking conditions were tested in random order: self-selected speed with 0% BWS (TM); self-selected speed with 30% BWS (BWS); and fastest comfortable speed with 30% BWS *and* nonparetic leg guidance (Guidance_NP_; described above). Walking trials were collected for as long as the participant could tolerate, up to a maximum of 40 seconds.

To isolate effects of TM, BWS, and Guidance_NP_, handrail hold was not provided and no AD or AFO’s were permitted during data collection. As with overground walking, an aircast was provided to control ankle instability if necessary.

#### Data Processing

Marker data were reduced using Vicon Nexus (Vicon Motion Systems Ltd., Version 1.6.1, Oxford, UK), modeled and filtered in Visual 3D (C-Motion, Version 4.82.0, Germantown, MD), and processed with custom Matlab (The MathWorks, Version 7.7.0 R2008b, Natick, MA) scripts to obtain spatiotemporal parameters for comparison with those obtained from the GAITRite^®^. Spatial measures were calculated using marker data and temporal variables were calculated from vertical ground reaction force data. Heel marker and ground reaction force data were filtered with a 4^th^ order bi-directional Butterworth lowpass filter (6Hz and 10Hz cutoff, respectively).

#### Statistical Analysis

Participant characteristics are provided in Table 1. Demographic data were analyzed to determine if differences existed between step length asymmetry groups using an α-level of 0.05 to identify significant differences. We used the Chi-Square test to determine if there were differences in side of paresis or sex among groups. To assess for differences in age and chronicity since stroke, we used the Kruskall-Wallis test.

**Table 1.** Demographics. Data for age, chronicity, and gait speeds are Mean ± SD. Data for LE Fugl-Meyer Synergy, Berg Balance Score, and Dynamic Gait Index are Median (Min, Max). *Abbreviations*: Symmetrical: paretic and nonparetic step lengths are equivalent; P_short_: paretic step length shorter than nonparetic; NP_short_: nonparetic step length shorter than paretic; yr: years; m/f: male/female; r/l: right/left; mo: months; m/s: meters per second Guidance_NP_: fastest comfortable walking speed, with 30% BWS, and nonparetic limb guidance; LE: lower extremity.

|  | **All** | **Symmetrical** | **P<sub>short</sub>** | **NP<sub>short</sub>** | **p-value** |
| --- | --- | --- | --- | --- | --- |
| n | 39 | 17 | 11 | 11 |  |
| age (yr) | 61.3 ± 11.4 | 63.4 ± 9 | 65.5 ± 8 | 53.9 ± 14.6 | 0.09 |
| sex (m/f) | 29/10 | 13/4 | 8/3 | 8/3 | 0.97 |
| paretic side (r/l) | 21/18 | 8/9 | 5/6 | 8/3 | 0.33 |
| chronicity (mo) | 68.4 ± 61.7 | 71.8 ± 68.5 | 48.3 ± 44.6 | 83.5 ± 65.9 | 0.38 |
| gait speed (m/s) |  |  |  |  |  |
| Overground (OG) | 0.63 ± 0.2 | 0.69 ± 0.2 | 0.59 ± 0.2 | 0.55 ± 0.16 | |
| Treadmill (TM) | 0.44 ± 0.14 | 0.48 ± 0.16 | 0.41 ± 0.12 | 0.4 ± 0.11 | |
| Body weight support (BWS) | 0.52 ± 0.2 | 0.59 ± 0.25 | 0.47 ± 0.14 | 0.47 ± 0.13 | |
| Guidance<sub>NP</sub> | 0.72 ± 0.25 | 0.78 ± 0.26 | 0.64 ± 0.18 | 0.72 ± 0.29 | |
| LE Fugl-Meyer Synergy (/22) | 16 (5, 22) | 16 (8, 22) | 20 (5, 22) | 13 (8, 20) | 0.1 |
| Berg Balance Score (/56) | 46 (32, 55) | 47 (40, 55) | 48 (41, 55) | 45 (32, 55) | 0.61 |
| Dynamic Gait Index (/24) | 15 (7, 22) | 15 (9, 22) | 12 (7, 21) | 15 (10, 19) | 0.81 |

Step length was investigated as the primary outcome. Secondary outcomes included: stride length and percent of gait cycle for single limb support (SLS%). Each variable was tested for normality. Separate 2-way ANCOVAs (Experimental Condition x Leg) were conducted for each asymmetry category to test for interaction effects among experimental conditions between paretic and nonparetic legs for step length and SLS%. A 1-way ANCOVA was performed on stride length to determine the effect of experimental condition for each asymmetry category. Gait speed was used as a covariate and retained in the respective model when found to be significant. In total, we performed 9 ANCOVAs; correcting for multiple comparisons, we established statistical significance at: α = 0.006 for all spatiotemporal variables. We used Tukey’s HSD to isolate differences when effects were detected. The Type I error rate was carried through and used for the respective post hoc analyses. All statistical tests were performed with JMP^®^ Pro (SAS Institute Inc. Version 13.2.0, Cary, NC) software.

## Results

### Overview

All participants (age: 61.3±11.4 yrs; 29 male; chronicity: 68.4±61.7 mo) experienced a single, monohemispheric stroke (confirmed with neuroimaging) and revealed hemiparesis, lower extremity (LE) motor dysfunction (median LE Fugl Meyer Synergy Score: 16/22, range: 5-22) and gait impairment (SSWS: 0.63±0.2 m/s; Table 1). The three step length asymmetry groups did not differ in demographic characteristics or clinical assessment of functional status (Table 1; all *p*’s>0.05). However, as described below, we identified differential patterns of response to the experimental conditions for each step length asymmetry subgroup (Table 2). All variables tested co-varied with gait speed (all *p*’s<0.0001).

**Table 2.** Asymmetry groups respond differently to experimental conditions. Data are mean ± SD. Reference values for single limb support duration (sec) are ≥ 0.67±0.03 sec for the overground walking speeds recorded in this study.^35^ *Abbreviations*: Symmetrical: paretic and nonparetic step lengths are equivalent; P_short_: paretic step length shorter than nonparetic; NP_short_: nonparetic step length shorter than paretic; OG: overground; TM: treadmill condition at self-selected walking speed, with 0% BWS; BWS: body weight support condition at self-selected walking speed, with 30% BWS; Guidance_NP_: fastest comfortable walking speed, with 30% BWS, and nonparetic limb guidance. * significant Experimental Condition X Leg interaction (*p*<0.0006) † significant Experimental Condition main effect (*p*<0.0006) ‡ significant Leg main effect (*p*<0.0006)

|  | Symmetrical |  | P<sub>short</sub> |  | NP<sub>short</sub> |  |
| --- | --- | --- | --- | --- | --- | --- |
|  | paretic | nonparetic | paretic | nonparetic | paretic | nonparetic |
| *Step length (m)* | * |  | ‡ |  | * |  |
| OG | 0.5 ± 0.09 | 0.5 ± 0.08 | 0.34 ± 0.12 | 0.47 ± 0.11 | 0.49 ± 0.08 | 0.37 ± 0.1 |
| TM | 0.35 ± 0.09 | 0.34 ± 0.09 | 0.2 ± 0.08 | 0.33 ± 0.09 | 0.35 ± 0.08 | 0.27 ± 0.13 |
| BWS | 0.41 ± 0.13 | 0.38 ± 0.12 | 0.28 ± 0.08 | 0.38 ± 0.09 | 0.4 ± 0.09 | 0.31 ± 0.11 |
| Guidance<sub>NP</sub> | 0.43 ± 0.15 | 0.52 ± 0.1 | 0.28 ± 0.13 | 0.48 ± 0.09 | 0.47 ± 0.13 | 0.48 ± 0.12 |
| *Single limb support (%)* | * |  |  |  | ‡ |  |
| OG | 0.26 ± 0.06 | 0.37 ± 0.05 | 0.25 ± 0.06 | 0.32 ± 0.05 | 0.23 ± 0.04 | 0.38 ± 0.06 |
| TM | 0.23 ± 0.06 | 0.29 ± 0.03 | 0.22 ± 0.05 | 0.26 ± 0.04 | 0.19 ± 0.06 | 0.33 ± 0.04 |
| BWS | 0.26 ± 0.05 | 0.3 ± 0.04 | 0.25 ± 0.04 | 0.27 ± 0.05 | 0.22 ± 0.05 | 0.34 ± 0.05 |
| Guidance<sub>NP</sub> | 0.31 ± 0.04 | 0.32 ± 0.05 | 0.3 ± 0.05 | 0.27 ± 0.05 | 0.28 ± 0.06 | 0.36 ± 0.03 |
| *Single limb support (sec)* | * |  |  |  | ‡ |  |
| OG | 0.38 ± 0.06 | 0.55 ± 0.12 | 0.35 ± 0.06 | 0.44 ± 0.07 | 0.37 ± 0.09 | 0.61 ± 0.09 |
| TM | 0.33 ± 0.1 | 0.43 ± 0.11 | 0.29 ± 0.09 | 0.34 ± 0.06 | 0.29 ± 0.11 | 0.50 ± 0.06 |
| BWS | 0.36 ± 0.08 | 0.43 ± 0.1 | 0.34 ± 0.03 | 0.37 ± 0.09 | 0.34 ± 0.12 | 0.52 ± 0.09 |
| Guidance<sub>NP</sub> | 0.41 ± 0.07 | 0.43 ± 0.08 | 0.38 ± 0.05 | 0.35 ± 0.08 | 0.39 ± 0.09 | 0.51 ± 0.09 |
| *Stride length (m)* | † |  | † |  |  |  |
| OG | 0.99 ± 0.17 | | 0.81 ± 0.22 | | 0.87 ± 0.17 | |
| TM | 0.68 ± 0.17 | | 0.54 ± 0.15 | | 0.62 ± 0.19 | |
| BWS | 0.79 ± 0.24 | | 0.66 ± 0.13 | | 0.71 ± 0.19 | |
| Guidance<sub>NP</sub> | 0.96 ± 0.23 | | 0.76 ± 0.18 | | 0.95 ± 0.25 | |

### Symmetrical step lengths (Symmetrical)

The Symmetrical group (n=17) was characterized by equivalent paretic (0.50±0.09m) and nonparetic (0.50±0.08) step lengths (PSR: 0.50 ± 0.01) while walking overground. We identified significant Experimental Condition x Leg interactions for step length and SLS% (*p*’s=0.0003). Step lengths were similar for the OG, TM, and BWS conditions while the nonparetic step length was longer during the Guidance_NP_ condition (Figure 1a,b). Importantly, relatively longer NP step length (0.52±0.1m) was achieved during Guidance_NP_ with a simultaneous reduction of paretic (0.43±0.15m) step length (Figure 1b). Even though the Symmetrical group participants exhibited spatial symmetry when walking overground, they revealed temporal asymmetry with a significantly reduced paretic SLS% (26±6%) relative to nonparetic SLS% (37±5%). Temporal asymmetry was noted during walking both on TM and with BWS (Figure 1c); however, the Guidance_NP_ condition induced temporal symmetry by increasing paretic SLS% (Δ: 5%) and decreasing nonparetic SLS% (Δ: 5%) concurrently (Figure 1d). Stride lengths achieved overground were reduced in the BWS and TM conditions but restored with the Guidance_NP_ condition (*p*=0.003; Figure 2).

**Figure 1.**
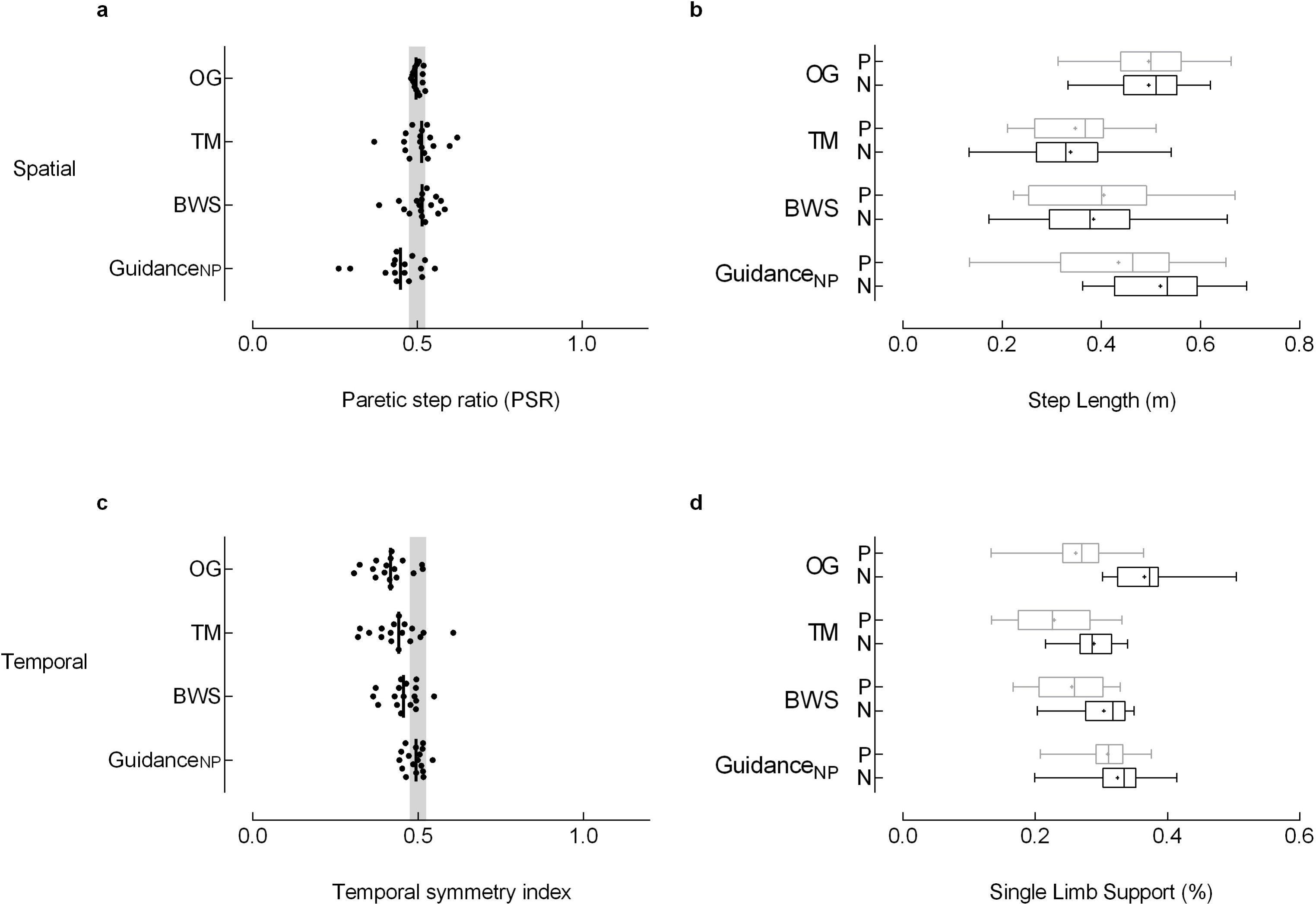
Symmetrical step lengths (Symmetrical) The Symmetrical group (n=17) was characterized by equivalent paretic and nonparetic step lengths while walking overground. **(a)** Spatial and **(c)** temporal symmetry was calculated with a symmetry index (SI) with the general equation SI =Xp/(Xp+Xnp), where Xp and Xnp are the paretic and nonparetic values for the variable of interest, respectively. Step length and percent of the gait cycle spent in single limb support were used to assess spatial and temporal symmetry. The symmetry index calculated for step length results in the paretic step ratio (PSR) used to categorize asymmetry groups (*see Methods*). Individual data are illustrated; the vertical black line represents the group median. The vertical gray shaded areas denote the SI values that represent symmetry (0.475 ≤ SI ≤ 0.525).^25^ Box-and-whisker plots for **(b)** step length and **(d)** single limb support duration (SLS%) illustrate the distribution of the individual leg data. The whiskers illustrate the 5^th^ and 95^th^ percentiles. Group means are depicted with “+”. Paretic and nonparetic leg data are illustrated in grey and black, respectively. Of note, the temporal symmetry achieved in the Symmetrical group with Guidance_NP_ results from a concurrent nonparetic reduction and paretic increase in SLS%. *Abbreviations*: OG: overground condition at self-selected walking speed; TM: treadmill condition at self-selected walking speed, with 0% BWS; BWS: body weight support condition at self-selected walking speed, with 30% BWS; Guidance_NP_: fastest comfortable walking speed, with 30% BWS, and nonparetic limb guidance.

**Figure 2.**
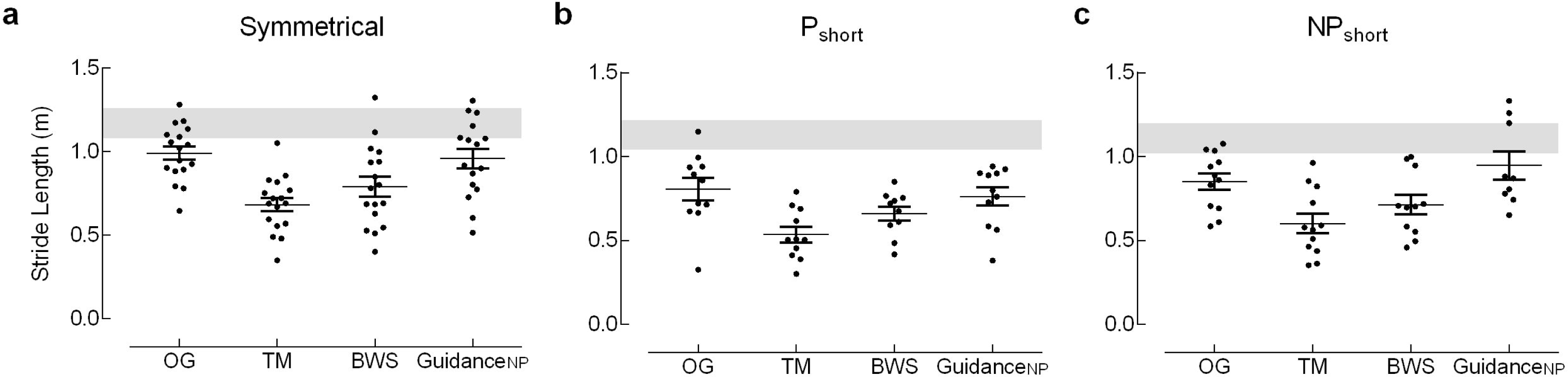
Stride length. Stride length depicts the combined length of the paretic and nonparetic steps. The shaded gray regions represent reference values (± 1 standard error) for overground stride length calculated from a known regression equation relating stride length and gait speed.^35^ Data are mean ± SEM. *Abbreviations:* Symmetrical: paretic and nonparetic step lengths are equivalent; P_short_: paretic step length shorter than nonparetic; NP_short_: nonparetic step length shorter than paretic; OG: overground condition at self-selected walking speed; TM: treadmill condition at self-selected walking speed, with 0% BWS; BWS: body weight support condition at self-selected walking speed, with 30% BWS; Guidance_NP_: fastest comfortable walking speed, with 30% BWS and nonparetic limb guidance.

### Paretic step length shorter than nonparetic (P_short_)

The P_short_ group (n=11) was characterized by a shorter paretic step length (0.34±0.12m) than nonparetic (0.47±0.11m) step length (PSR: 0.41 ± 0.05) while walking overground. While we did not find a significant Experimental Condition x Leg interaction for step length (*p*=0.07), we did identify a main effect of Leg; the shorter paretic step was consistent across all walking conditions (*p*=0.0001; Figure 3a,b). We identified a significant main effect for Experimental Condition for stride length (*p*=0.005); BWS produced stride lengths similar to OG while the TM and Guidance_NP_ conditions produced stride lengths less than OG. We identified a tendency for the Experimental Condition x Leg interaction for SLS% (*p*=0.006); while walking overground, nonparetic SLS% (32±5%) was greater than paretic (25±6%), but SLS% approached symmetry in the TM, BWS, and Guidance_NP_ conditions though these changes failed to reach statistical significance given the adjusted ±-level. Indeed, the median of the symmetry index for SLS% fell within the range of symmetry for the Guidance_NP_ condition (Figure 3c). During Guidance_NP_, we observed a concurrent decrease in nonparetic SLS% (Δ: 5%) and increase in paretic SLS% (Δ: 5%; Figure 3d).

**Figure 3.**
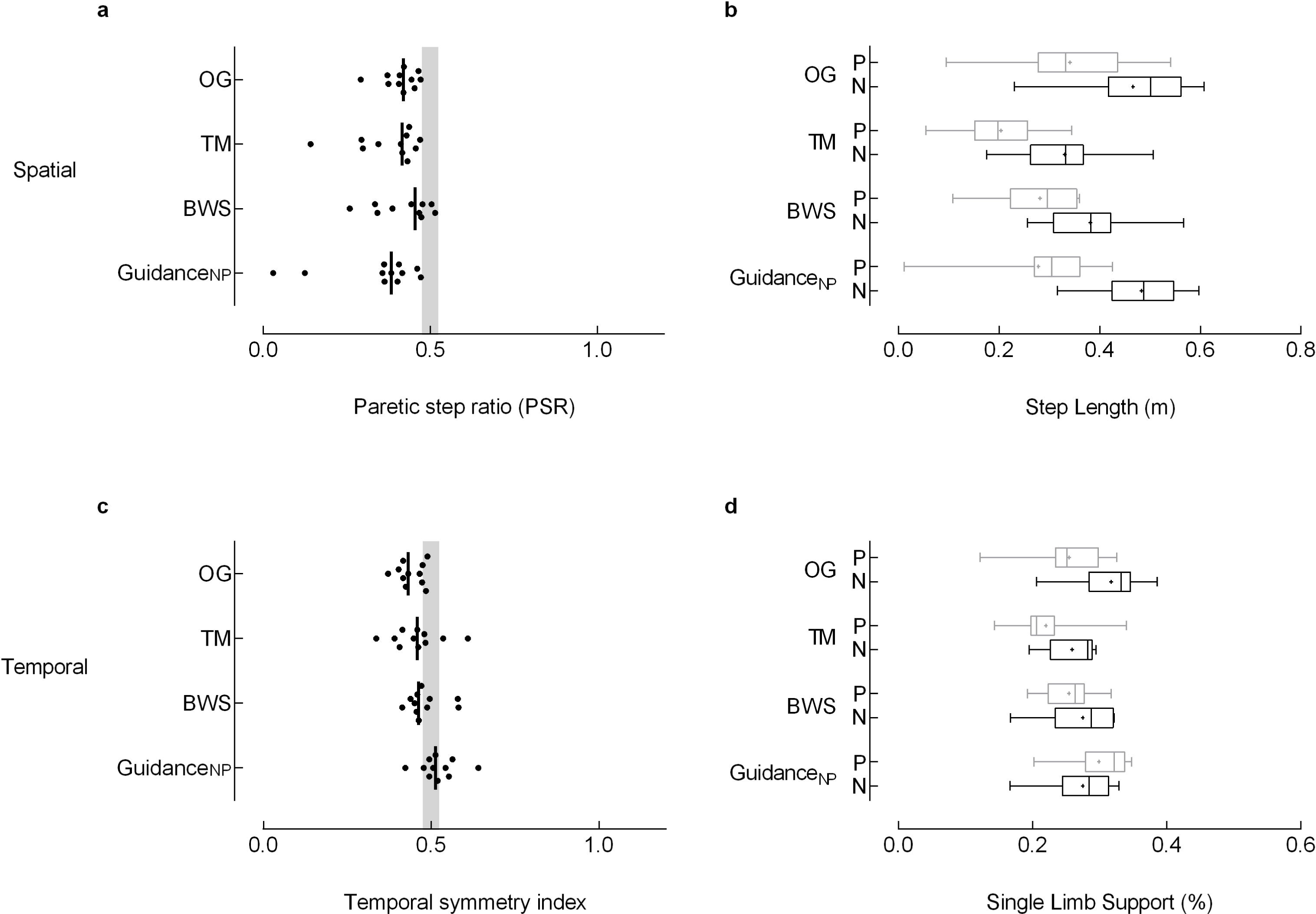
Paretic step length shorter than nonparetic (P_short_) The P_short_ group (n=11) was characterized by a shorter paretic step length than nonparetic step length while walking overground. Spatial (top, left) and temporal (bottom, left) symmetry was calculated with a symmetry index (SI) with the general equation SI =Xp/(Xp+Xnp), where Xp and Xnp are the paretic and nonparetic values for the variable of interest, respectively. Individual data are illustrated; the vertical black line represents the group median. The vertical gray shaded areas denote the SI values that represent symmetry (0.475 ≤ SI ≤ 0.525).^25^ Box-and-whisker plots for step length (top, right) and single limb support duration (SLS%; bottom, right) illustrate the distribution of the individual leg data. The whiskers illustrate the 5^th^ and 95^th^ percentiles. Group means are depicted with “+”. Paretic and nonparetic leg data are illustrated in grey and black, respectively. Note, a concurrent decrease in nonparetic SLS% (Δ: 5%) and increase in paretic SLS% (Δ: 5%) between the overground and Guidance_NP_ conditions (c). While these changes resulted in temporal symmetry, they failed to reach statistical significance. *Abbreviations*: OG: overground condition at self-selected walking speed; TM: treadmill condition at self-selected walking speed, with 0% BWS; BWS: body weight support condition at self-selected walking speed, with 30% BWS; Guidance_NP_: fastest comfortable walking speed, with 30% BWS, and nonparetic limb guidance; P: paretic; N: nonparetic.

### Nonparetic step length shorter than paretic (NP_short_)

The NP_short_ group (n=11) walked with shorter nonparetic (0.37±0.01m) than paretic step (0.49±0.08m) lengths (PSR: 0.58 ± 0.05) overground. We identified a significant Experimental Condition x Leg interaction for step length (*p*=0.001). While the spatial asymmetry noted OG was present in the TM and BWS conditions, the Guidance_NP_ condition produced symmetric step lengths in the NP_short_ group (Figure 4a) by increasing the nonparetic step length (Δ: 0.11m; Figure 4b). A main effect of Leg (*p*<0.0001) confirmed paretic SLS% less than nonparetic across all walking conditions; no interaction effects were detected for SLS% (Figure 4 c,d). Importantly, P SLS% increased from 23% to 28% of the gait cycle between OG and Guidance_NP_ conditions, though this increase did not achieve statistical significance.

**Figure 4.**
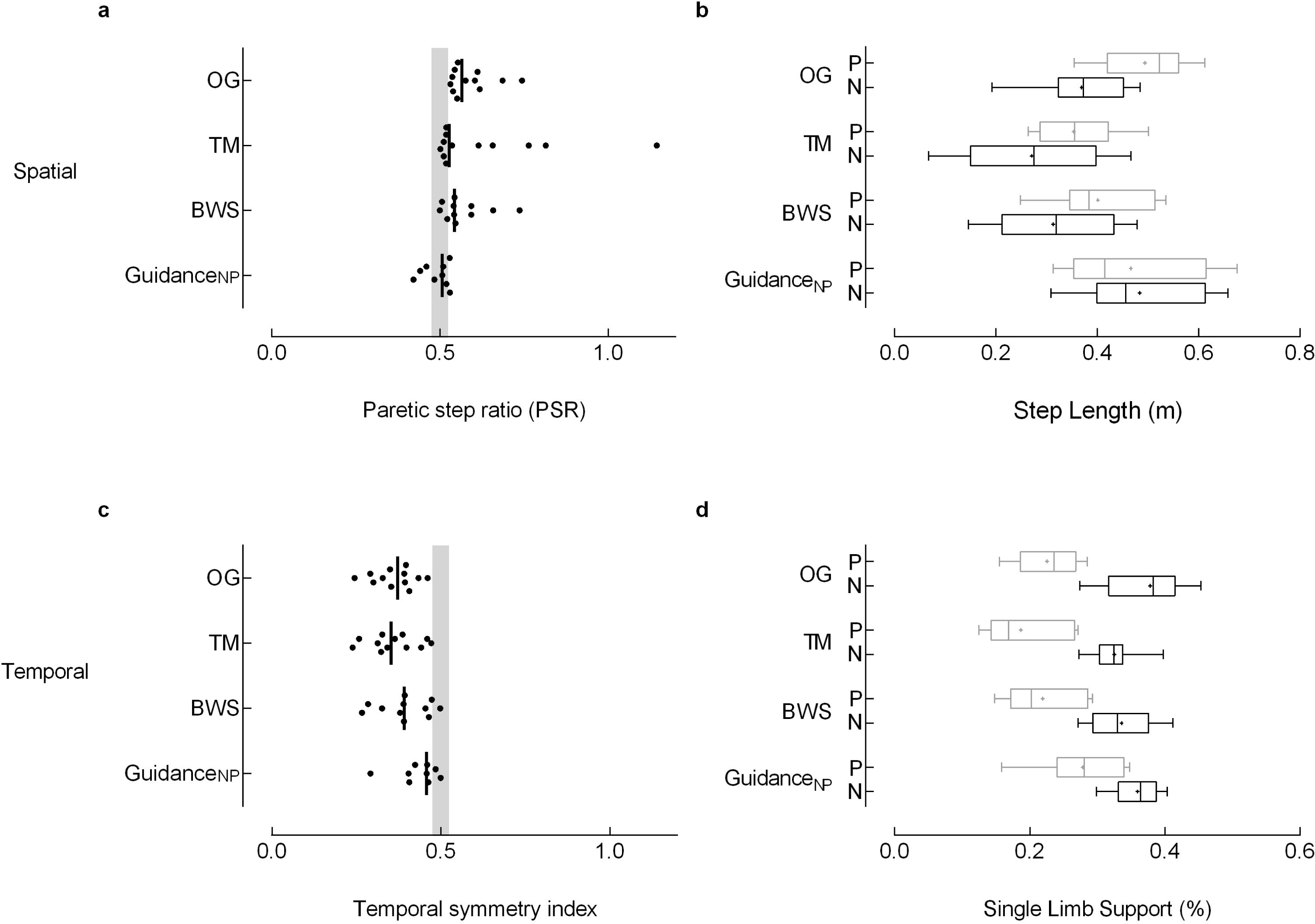
Nonparetic step length shorter than paretic (NP_short_) The NP_short_ group (n=11) walked with shorter nonparetic than paretic step lengths overground. Spatial (top, left) and temporal (bottom, left) symmetry was calculated with a symmetry index (SI) with the general equation SI =Xp/(Xp+Xnp), where Xp and Xnp are the paretic and nonparetic values for the variable of interest, respectively. Individual data are illustrated; the vertical black line represents the group median. The vertical gray shaded areas denote the SI values that represent symmetry (0.475 ≤ SI ≤ 0.525).^25^ Box-and-whisker plots for step length (top, right) and single limb support duration (SLS%; bottom, right) illustrate the distribution of the individual leg data. The whiskers illustrate the 5^th^ and 95^th^ percentiles. Group means are depicted with “+”. Paretic and nonparetic leg data are illustrated in grey and black, respectively. Note, the Guidance_NP_ condition produced symmetric step lengths (a) by increasing the nonparetic step length (b; Δ: 0.11m). Importantly, P SLS% increased from 23% to 28% of the gait cycle between OG and Guidance_NP_ conditions (d), though this increase did not achieve statistical significance. *Abbreviations*: OG: overground condition at self-selected walking speed; TM: treadmill condition at self-selected walking speed, with 0% BWS; BWS: body weight support condition at self-selected walking speed, with 30% BWS; Guidance_NP_: fastest comfortable walking speed, with 30% BWS, and nonparetic limb guidance; P: paretic; N: nonparetic.

## Discussion

Here we investigated whether groups with different spatial asymmetry patterns responded uniquely to task conditions that comprise components of the LT experience. Based on task condition we detected differential responses in step length and single limb support percentage across asymmetry groups. We expected the Guidance_NP_ condition would: 1) increase the nonparetic step length, regardless of group, and 2) normalize temporal asymmetry, especially in the NP_short_ group. However, our results indicate we increased nonparetic step length for only the NP_short_ group. Furthermore, while Guidance_NP_ improved temporal symmetry for the Symmetrical group, temporal asymmetry persisted in the NP_short_ and P_short_ groups.

### Effects induced by locomotor training parameters

Each seemingly simple decision regarding the training environment and parameters can influence patient response. Indeed, prior work reported improvement of single limb support symmetry simply by walking on the treadmill or using body weight support.^16,21,31^ Our results contrast with these findings. However, the improved symmetry during treadmill walking noted in previous studies occurred simultaneously with use of handrails^16,31^ and may, therefore, be an artefact of increased postural support available through upper extremity support rather than a direct response to a treadmill-induced perturbation.^21,22^ Generally, we found the treadmill and use of body weight support insufficient to induce either spatial or temporal symmetry. The Guidance_NP_ condition was more successful in inducing symmetry, although responses differed by asymmetry subgroup.

### Effects in the context of asymmetry subgroups

The underlying premise that a single task-specific training approach would positively benefit a group with heterogeneous gait deficits limits opportunity to better understand the interaction between subgroup characteristics and treatment effects. Interestingly, in relatively homogenous samples intentionally represented by people with a short nonparetic relative to paretic step length, improved spatial symmetry was noted in response to each of two different training paradigms (i.e., split-belt training and unilateral step training).^18,32^ The Guidance_NP_ condition was similarly able to induce spatial symmetry in the NP_short_ group. However, spatial symmetry was not achieved in other subgroups during the Guidance_NP_ condition.

From the literature, we also note that improvements in temporal symmetry appear more elusive.^18,32^ For example, in a case series, Lewek reported improved temporal symmetry in an individual who started with spatial symmetry and temporal asymmetry; however, in an individual who started with both spatial and temporal asymmetries, the temporal asymmetry remained unchanged despite improvements in spatial symmetry.^33^ Though we did not investigate treatment response (i.e., exposure to repeated sessions) in the current study, our findings align with prior work.^18,32,33^ Guidance_NP_ was able to induce temporal symmetry acutely only in the Symmetrical group.

### Is targeting improved symmetry sufficient?

Recent studies have reported improved gait symmetry as an acute effect^16,31^ or a treatment outcome of gait-related interventions.^12,18,32,33^ While others might interpret an improved symmetry ratio as a positive effect, we argue that symmetry ratios can be misleading. A change in symmetry ratio alone cannot elucidate the source of change, specifically, whether improvements result from changes in the paretic, nonparetic, or both limbs.^12,31,33^ Improved symmetry ratios alone are therefore insufficient to conclude a beneficial outcome has occurred. Indeed, when individual leg changes are reported, the data illustrate that improved symmetry ratios are often achieved through a non-physiologic reduction from the nonparetic limb with no, or only nominal, improvement noted in the paretic limb.^16,18^ Consistent with prior work, we observed a decrease in nonparetic SLS% on the TM (not tested explicitly; Table 2) across all groups.^16^ While it could be argued that this change produced improved symmetry (e.g., through reduction of the between-leg difference^16^), the changes induced on the TM did not achieve our definition of symmetry, neither did they approach physiologic durations of paretic single limb support. Of note, our externally-guided condition, Guidance_NP_, was the only experimental condition that restored single limb support symmetry between legs. However, this restoration of temporal symmetry was only observed in the spatially symmetrical group. Furthermore, the change in SLS% symmetry during Guidance_NP_ was driven by concurrent changes in both legs: an increase in paretic single support and a concomitant reduction in nonparetic single support. Notably, these changes were consistent with a physiologic gait pattern used by healthy controls walking on a treadmill, which is characterized by single limb support duration of ∼31-32% of the gait cycle.^34^ We emphasize these observations were made acutely, in response to an experimental condition.

## Conclusions/Implications

Commonly used rehabilitation interventions for gait dysfunction following stroke do not produce uniform effects. We identified differential acute responses to locomotor training conditions between groups with disparate asymmetry patterns, suggesting these subgroups may benefit from distinct intervention strategies. Improvements in temporal symmetry revealed in the Symmetrical group were noted to result from both limbs. Similarly, improvements in spatial symmetry noted in the NP_short_ group were driven by bilateral improvements, namely increased nonparetic step length occurring in combination with increased paretic single limb support. By investigating individual limb effects, we were able to determine the changes in spatial and temporal symmetry resulted from desirable effects rather than compensatory mechanisms.

## Acknowledgements

This research was supported by the Department of Veterans Affairs, Rehabilitation Research & Development Service, Project #A6365B (SAK &CP) and Research Career Scientist Awards #F7823S (CP) and #A9272S (SAK) and the VA Brain Rehabilitation Research Center of Excellence (B6793C). Dr. Little received support from the Foundation for Physical Therapy and NIH T32 Neuromuscular Plasticity Training Grant (No. 5 T32 HD043730-08, K Vandenborne, PI). Dr. Mercado received support from the University Scholars’ Program at the University of Florida. Dr. Kautz also received support from NIH P20 GM109040.

This material is the result of work supported with resources and the use of facilities at the NF/SG Veterans Administration Health Care System, Gainesville, FL, USA. The contents do not represent the views of the Department of Veterans Affairs, the NIH, or the United States Government. The funding source played no role in either writing this manuscript or the decision to submit for publication. The corresponding author retains full access to all data in the study and assumes final responsibility for the decision to submit for publication.

We thank: Helen Emery for her assistance with data collection and Yueh-Yun Chi, PhD for her assistance with the statistical analysis.

## Footnotes

† The requisite sensory inputs underscoring locomotor training include: 1) afferent stimuli from limb loading, 2) proper trunk alignment and upright orientation relative to gravity, 3) hip extension, 4) appropriate walking speed, and 5) phasic timing of loading and unloading cycles during walking.8

